# A draft genome of *Alliaria petiolata* (garlic mustard) as a model system for invasion genetics

**DOI:** 10.1101/2021.02.17.431678

**Authors:** Nikolay Alabi, Yihan Wu, Oliver Bossdorf, Loren H. Rieseberg, Robert I. Colautti

## Abstract

The emerging field of invasion genetics examines the genetic causes and consequences of biological invasions, but few study systems are available that integrate deep ecological knowledge with genomic tools. Here we report on the *de novo* assembly and annotation of a genome for the biennial herb *Alliaria petiolata* (M. Bieb.) Cavara & Grande (Brassicaceae), which is widespread in Eurasia and invasive across much of temperate North America. Our goal was to sequence and annotate a genome to complement resources available from hundreds of published ecological studies, a global field survey, and hundreds of genetic lines maintained in Germany and Canada. We sequenced a genotype (EFCC-3-20) collected from the native range near Venice, Italy and sequenced paired-end and mate pair libraries at ~70 × coverage. A *de novo* assembly resulted in a highly continuous draft genome (N_50_ = 121 Mb; L_50_ = 2) with 99.7 % of the 1.1 Gb genome mapping to scaffolds of at least 50 Kb in length. A total of 64,770 predicted genes in the annotated genome include 99 % of plant BUSCO genes and 98 % of transcriptome reads. Consistent with previous reports of (auto)hexaploidy in western Europe, we found that almost one third of BUSCO genes (390/1440) mapped to two or more scaffolds despite < 2 % genome-wide average heterozygosity. The continuity and gene space quality of our draft assembly will enable molecular and functional genomic studies of *A. petiolata* to address questions relevant to invasion genetics and conservation strategies.

## Introduction

Biological invasions are a threat to global biodiversity with significant impacts to human health and welfare (Mack et al., 2000). They also present opportunities for large-scale ‘natural’ experiments to study ecological and evolutionary processes in the wild (Mooney and Cleland, 2001; Sax et al., 2005). Despite a large body of ecological research and a growing number of evolutionary studies of invasive species, functional genetic and ‘omics approaches have been rare among studies of invasive species until recently (reviewed in (Barrett, 2015; Bock et al., 2015); but see e.g. (Barrett et al., 2016; Boheemen et al., 2017; Bourne et al., 2020)). Apart from studies using neutral markers to assess population structure (reviewed in (Dlugosch and Parker, 2008)), relatively little is known about the genetic causes and consequences of biotic invasions at the molecular level. Perhaps this is partly due to a lack of genomic resources, which has hindered high-resolution genetic studies of most invasive species. Here, we report on a draft genome for the herbaceous biennial plant *Alliaria petiolata,* a plant invader in North America with potential to become a model system for the emerging field of invasion genetics.

Several factors favour *A. petiolata* as an emerging model system for invasion genetics (Colautti et al., 2014). First, it is widely distributed with variable ecological impacts throughout its range (Lankau et al., 2009; USDA, 2020). Second, it has a relatively simple two-year life cycle with well-defined life stages (seed, rosette, bolting, senescent). This simple life history facilitates measurements of lifetime fitness, natural selection, and their combined influence on population dynamics in natural populations. Third, *A. petiolata* is a member of the Brassicaceae and therefore benefits from genetic resources available for well-studied species like *Brassica rapa* (canola) and the model plant *Arabidopsis thaliana*, providing opportunities for functional and comparative genomics. Fourth, high selfing rates produce naturally inbred seed families that can be maintained through single-seed descent. Fifth, *A. petiolata* has been the focal species in hundreds of field surveys and experimental studies, including influential studies testing the role of natural enemies (Lewis et al., 2006), competition (Prati and Bossdorf, 2004), the ‘novel weapons’ hypothesis (Callaway et al., 2008), competitive ability (Bossdorf et al., 2004), glucosinolate metabolism (Haribal et al., 2001), and eco-evolutionary dynamics (Lankau et al., 2009). One mechanism that is especially well-studied is the production of secondary metabolites and their effects on soil microbiota, particularly the suppression of mycorrhizal fungi that form beneficial symbiotic networks among native plant roots (Anthony et al., 2017; Duchesneau et al., 2021). Understanding the genomic basis of such interactions with soil microbes will not only advance basic science but it also has potential applications in plant restoration and agriculture.

Recent efforts to develop *A. petiolata* as a model system include the Global Garlic Mustard Field Survey (GGMFS), which mobilized 164 participants from 16 countries to collect field data and seed samples across Europe and North America, resulting in thousands of seed families from 383 distinct populations (Colautti et al., 2014). A subset of inbred lines collected across North America have been maintained through single-line descent in labs in Germany (Bossdorf Lab, University of Tübingen) and Canada (Colautti Lab, Queen’s University). Adding to these resources, we here report on a draft genome of a single *A. petiolata* genotype from Europe, annotated with RNA sequencing of leaf and root tissue.

## Methods & Materials

### Study Organism and Line Derivation

*Alliaria petiolata* (M. Bieb.) Cavara & Grande is a biennial herbaceous plant in the Thlaspideae tribe of the Brassicaceae family. It was introduced to North America prior to 1868, when it was discovered in Long Island, New York (Nuzzo, 1993). By 1948 it was reported on the West Coast of North America and has established in at least 37 U.S. States and five Canadian provinces from the Atlantic coast to the Pacific (Cavers et al., 1979; USDA, 2020). As the only species of *Alliaria* with a broad distribution, *A. petiolata* is relatively easy to identify in natural habitats owing to its white flowers and dentated peltate leaves with long petioles. It is considered a noxious weed across most of its introduced range, due in part to impacts on native plants and tree regeneration in deciduous forest ecosystems (Cipollini and Cipollini, 2016; Stinson et al., 2007). Two genome types have been identified, including diploids (2*n* = 14) in Eastern Europe and Western Asia and hexaploids (2*n* = 42 and 2*n* = 36) in Central Europe and North America (Esmailbegi et al., 2018; Weiss-Schneeweiss and Schneeweiss, 2003). The haploid genome size was previously estimated to be 1.35 Gb based on flow cytometry (Barow and Meister, 2002).

#### S_0_ generation

All source material for genome and transcriptome sequencing originated from a single individual grown from seed collected as part of the GGMFS (Colautti et al., 2014). The inbred line used in this study is from population EFCC3, collected in 2011 from a small forest fragment (~2,000 m^2^) surrounded by agricultural land, about 75 km northwest of Venice, Italy (UTM 45.71 °N × 11.72 °E). The specific seed line was sampled at 3.2 m along a 10 m sampling transect originating at the edge of population EFCC3. This inbred line (code EFCC3-3-20) is currently maintained along with other GGMFS seed collections by two of the coauthors in replicate collections in Tübingen, Germany (Bossdorf) and Kingston, Ontario, Canada (Colautti).

#### S_1_ generation

In July 2012, ten seeds of the EFCC3-3-20 genotype from the original S_0_ field collection were removed from cold storage (4 °C), surface washed with a mild detergent and rinsed with distilled H_2_O before surface sterilizing in 10 % bleach for 10 minutes. Sterilized seeds were again rinsed with distilled H_2_O before placing on filter paper saturated with distilled H_2_O and sealed in a petri dish with paraffin wax. We stratified seeds in the dark at 10 °C for ~ 90 days and thereafter inspected weekly until emerging radicles were observed. Germinating seeds were transplanted into 4” plastic pots containing a peat soil mixed with vermiculite that was watered to saturation and placed under shade cloth in the Horticulture Greenhouses at the University of British Columbia. We let seedlings establish in soil, watered as needed, for four weeks before a small amount (~ 5 mm x 2 mm) of young leaf meristem tissue was harvested from a single individual and immediately preserved in liquid nitrogen. Roughly 25 to 50 mm^3^ of this tissue was divided into two separate 2 mL screw-cap tubes, each containing two stainless steel ball bearings of 2mm diameter. These samples were used for genomic DNA purification and genome sequencing.

A second individual from the same inbred family was transplanted to an 8” plastic pot and fertilized with 20/20/20 N/P/K fertilizer to encourage rosette growth before being moved outside for cold vernalization from October 2012 to April 2013 at the University of British Columbia Horticulture Greenhouses. In April 2013, we moved the plant back into the greenhouse and sprayed with 2 % insecticidal soap to remove pests. Once inside the greenhouse, the plant was left to mature and set seed autonomously via self-pollination. Mature siliques were harvested in July 2013 and seeds were stored in paper envelopes at 4 °C.

#### S_2_ generation

In May 2016, we removed a subset of 10 seeds of the S_1_ generation from cold storage and germinated in a 60 mm × 15 mm petri dish containing filter paper covered with a mixture of autoclaved ProMix soil and silica sand (1:9 ratio). We added distilled water until saturation and thereafter petri dishes were sealed with paraffin wax before storing in the dark at 4 °C. Of these, six seedlings germinated and were retained for transcriptome sequencing, divided into one of two treatments. The first true leaf from each of the three plants in the experimental treatment were cut with scissors. We used a Kimwipe tissue saturated with either 0.4 mM jasmonic acid (JA) dissolved in 10 % ethanol (treatment) or 10 % ethanol alone (control), adhered directly to maintain contact the cut site (treatment) or uncut leaf (control). We replaced the saturated Kimwipe every 8 h to maintain the signal. After 48 h of treatment, we harvested seedlings and preserved treated leaves and root tissue in liquid nitrogen, to be used for RNA purification and transcriptome sequencing.

### DNA Isolation & Library Construction

In September 2013, frozen tissue from the S_1_ genotype was pulverized and extracted using a Cetyltrimethyl Ammonium Bromide (CTAB) protocol (Clarke, 2009) with the following modifications. After pulverising tissue in a bead mill homogenizer at 60 Hz for 60 s, we added 1 mL chilled wash buffer (200 mM Tris-HCl pH 8.0, 50 mM EDTA, 250 mM NaCl) and incubated on ice for 10 min. The purpose of this wash step is to remove secondary metabolites after disruption of cell walls but prior to cell lysis with CTAB. Following this initial wash step, we spun tubes in a microcentrifuge at 4000 g and 4° C for 10 minutes, then discarded the supernatant and added another 1 mL of wash buffer. This was repeated once more for a total of three wash cycles until no coloration was visible in the supernatant. After final discard of the supernatant and addition of warm lysis buffer as per the CTAB protocol, we vortexed tubes briefly to resuspend plant cells. After completion of the CTAB protocol, pellets were dissolved in 50 μL of reverse osmosis (RO) H_2_O and sent to Centre d’expertise et de services Génome Québec (Génome Québec) for library preparation and sequencing.

We used four separate sequencing libraries for genome assembly: (i) One whole-genome shotgun sequencing library using the Illumina TruSeq DNA v1 preparation kit with a target fragment length of 150 bp. (ii & iii) Two Illumina Nextera MatePair libraries with target lengths of 5 Kb and 10 Kb. These three libraries were multiplexed and sequenced on a single flowcell of Illumina HiSeq 2000 using 2 × 100 bp paired end (PE) sequencing chemistry. (iv) Target fragment lengths of 450 bp using the Illumina TruSeq DNA v1 and sequenced on Illumina MiSeq with 2 × 250 bp paired-end reads.

### RNA Isolation & Library Construction

For RNA purification, we pulverized frozen leaf and root tissue In March 2017, in the same manner as the DNA extraction protocol outlined in the previous section. After pulverizing the tissue, we extracted whole RNA from each plant separately using Invitrogen’s TRIzol reagent, following the manufacturer’s protocol (Pub No. MAN0001271 Rev. A.0). We sequenced four of the six extractions with the highest RNA yields at Génome Québec using the KAPA rRNA-depletion Kit (plant) followed by the Illumina TruSeq LT multiplex kit for sequencing on a single lane of Illumina HiSeq 2500 with 2 × 125 bp paired-end reads. Eight separate libraries were sequenced based on tissue and treatment as described above (see S2 Generation): Control Leaf (CL), Control Root (CR), Treated Leaf (TL), and Treated Root (TR), with two replicates each.

### Data Processing Methods

Raw sequencing data was processed and demultiplexed by Génome Québec, and copied to the Frontenac cluster hosted by the Centre for Advanced Computing (CAC) at Queen’s University and the Cedar cluster maintained by Simon Fraser University on behalf of Compute Canada. The CAC maintains the Rosalind Franklin Cluster for Analysis of Complex Genomes, which is a 256-core computing cluster with 2 TB of RAM. We used this hardware for the memory-intensive steps of *de novo* genome assembly, with the remaining analyses completed using shared Frontenac and Cedar clusters, and on personal computers. All FASTQ files from both experiments passed quality controls using **fastqc** (version 0.11.5) (Andrews, 2010). We used the raw, demultiplexed FASTQ files for *de novo* assembly, but the transcriptome data were pre-processing using **cutadapt** (Martin, 2011) to trim adapters and remove reads shorter than 25b prior to assembly.

Our genome assembly pipeline involved two main steps. First, we used **ALLPATHS-LG** (Gnerre et al., 2011) version R52488 to assemble contigs from both the HiSeq and MiSeq paired-end libraries and then to link contigs into scaffolds using the 5 Kb and 10 Kb MatePair libraries. We filtered reads that aligned to a draft cpDNA genome but otherwise did not pre-process reads as the **ALLPATHS-LG** pipeline includes filtering steps. Mitochondrial DNA sequences were manually removed from the final assembly. The analysis parameters included PLOIDY = 2 and HAPLOIDIFY = TRUE to perform a diploid genome assembly. Although our genome is likely hexaploid, polyploid models are not supported by **ALLPATHS-LG** and low heterozygosity is expected given the high selfing rates in natural populations. Second, we joined scaffolds from **ALLPATHS-LG** into larger mega-scaffolds using **redundans** (Pryszcz and Gabaldón, 2016) version 0.13c, with the following parameters: *identity* = 0.9, *iters* = 5, *joins* = 5, *limit* = 1, *linkratio* = 0.7, *mapq* = 10, *minlength* = 1000 and *overlap* = 0.75. Thus, overlapping scaffolds were collapsed only if they overlapped by at least 75% with an average identity of at least 90%. We repeated this script four times with output scaffolds of the prior run acting as input scaffolds given the long run-time required (~28d). This second round of assembly combines scaffolds with overlapping similarity, resulting in mega-scaffolds that can span across multiple chromosomes, and other regions of high recombination.

Sequences from the transcriptome experiment were cleaned and assembled with **trinity** (Grabherr et al., 2011) following protocols outlined on the software documentation and in Haas *et al* (Haas et al., 2013). We used default parameters and the quality of the assembly was analyzed using the custom perl scripts included in the **trinity** package to examine full length transcripts and scaffolds (i.e. Contig Nx lengths in *analyze_blastPlus_topHit_coverage.pl, TrinityStats.pl*). Additionally, we mapped read pairs to the transcriptome assembly to assess read content using **bowtie2** (version 2.3.3.1) with default parameters (Langmead and Salzberg, 2012). We used **TransDecoder** to predict open reading frames in transcripts before using **Trinotate** to annotate and analyze assembled transcripts (Haas et al., 2013).

As a first step in annotation, we established a detailed repeat library. Miniature Inverted Transposable Elements (MITES) were identified using **MITE Tracker** (Crescente et al., 2018). Long Terminal Repeat (LTR) elements were identified using the **GenomeTools** package (Gremme et al., 2013). To reduce the number of false positive LTR transposons, only those that contained PPT (poly purine extract) or PBS (primer binding sites) within a 30bp radius were kept and the rest filtered. We further filtered the LTR candidates to eliminate three main sources of false positives: tandem local repeats such as centromeric repeats, local gene clusters derived from recent gene duplications, and two other transposable elements located in adjacent regions. We also identified elements with nested insertions. After processing known MITEs and LTR elements, we identified additional repetitive sequences using **RepeatModeler** against a transposase database and excluded gene fragments using **ProtExcluder**.

The annotation pipeline **MAKER** (Campbell et al. 2014) was used to identify gene models and predict functional annotations in the draft genome. Both *est2genome* and *protein2genome* modes were used initially to make *ab initio* gene predictions from EST and protein evidence, respectively. The EST evidence was based on our own transcriptome data whereas the protein evidence was gathered from the reference proteomes of six closely related plants available from the SwissProt database: *Arabidopsis thaliana, Glycine max, Brassica oleracea, Medicago truncatula, Brassica napus,* and *Brassica rapa*. From the first round of annotation, high confidence models were predicted by the *maker2zff* command with default minimums (50 % of splice sites confirmed by EST alignment, 50 % of exons match an EST alignment, 50 % of exons overlap any evidence, and maximum AED of 0.5). These predictions were used to train the *ab initio* gene predictor **SNAP** (Korf, 2004). A second round of **MAKER** was run using the hidden Markov model (HMM) from **SNAP** rather than the *est2genome* mode. All other settings were the same as for the first run, with the transcripts now being used only as evidence to support *ab initio* gene predictions. Two more rounds of annotation and gene prediction improvement followed. An **Augustus** gene prediction file was also generated for use as a second *ab initio* prediction (Stanke and Morgenstern, 2005). For the final round of annotation in **MAKER**, the final HMM file from **SNAP**, the *A. petiolata* species **Augustus** library and gene prediction file were all used in addition to the following settings: always complete, single exons, and using correct EST fusion.

We used Benchmarking Universal Single-Copy Orthologs (**BUSCO**) v4.1 (Simão et al., 2015) to assess the assembly quality of the final draft genome. We used plant lineage delineation from the EmbryophytaDB V10 database, focusing on universal orthologs present in > 90 % of lineages, resulting in a total of 1,440 BUSCO orthologs. We used **minimap2** to check for assembly contiguity and synteny with the model plant *Arabidopsis thaliana* using the TAIR 10 assembly and up to 20 % sequence divergence. We used the R package ***pafr*** to generate dot-plots for the four largest scaffolds of *A. petiolata* and all chromosomes of *A. thaliana*.

The genome annotation files were curated through **deFusion** to resolve fused gene annotation problems (Wang et al., 2021). We also used **EvidenceModeler** (Haas et al., 2008) to combine *ab initio* gene predictions and protein and transcript alignments into weighted consensus gene structures. The functional annotations were then created using NCBI **BLAST+** and **InterProScan** (Jones et al., 2014) by adding new names, domains, and putative functions to improve the utility of the genome database.

### Data Availability

Raw data used for genome assembly, transcriptome assembly, and the final draft genome is available from DDBJ/ENA/GenBank under BioProject **PRJNA702530**, including the Whole Genome Shotgun project **AHZTW000000000** from **BioSample SAMN17958863**, with sequence reads deposited in **SRR13765007 - SRR13765010**. The transcriptome analysis BioSamples **SAMN17984654 - SAMN17984657** and raw sequence reads are also available (**SRR13765003 - SRR13765006**). The latest genome assembly scripts are available on GitHub (https://github.com/ColauttiLab/Ap_Genome_Assembly).

## Results and Discussion

Most (98.7 %) of the essential single-copy genes from **BUSCO** mapped to our assembly (Fig. 1), with 71.5 % occurring only once in the assembled genome. Similarly, 98 % of sequence reads from the transcriptome experiment mapped successfully to the assembled genome. Sequencing of the transcriptome libraries yielded a total of 68.1 Gb from 272.5 million paired reads. Trimming sequence reads for quality reduced usable data by less than 2 %.

**Figure 1.**
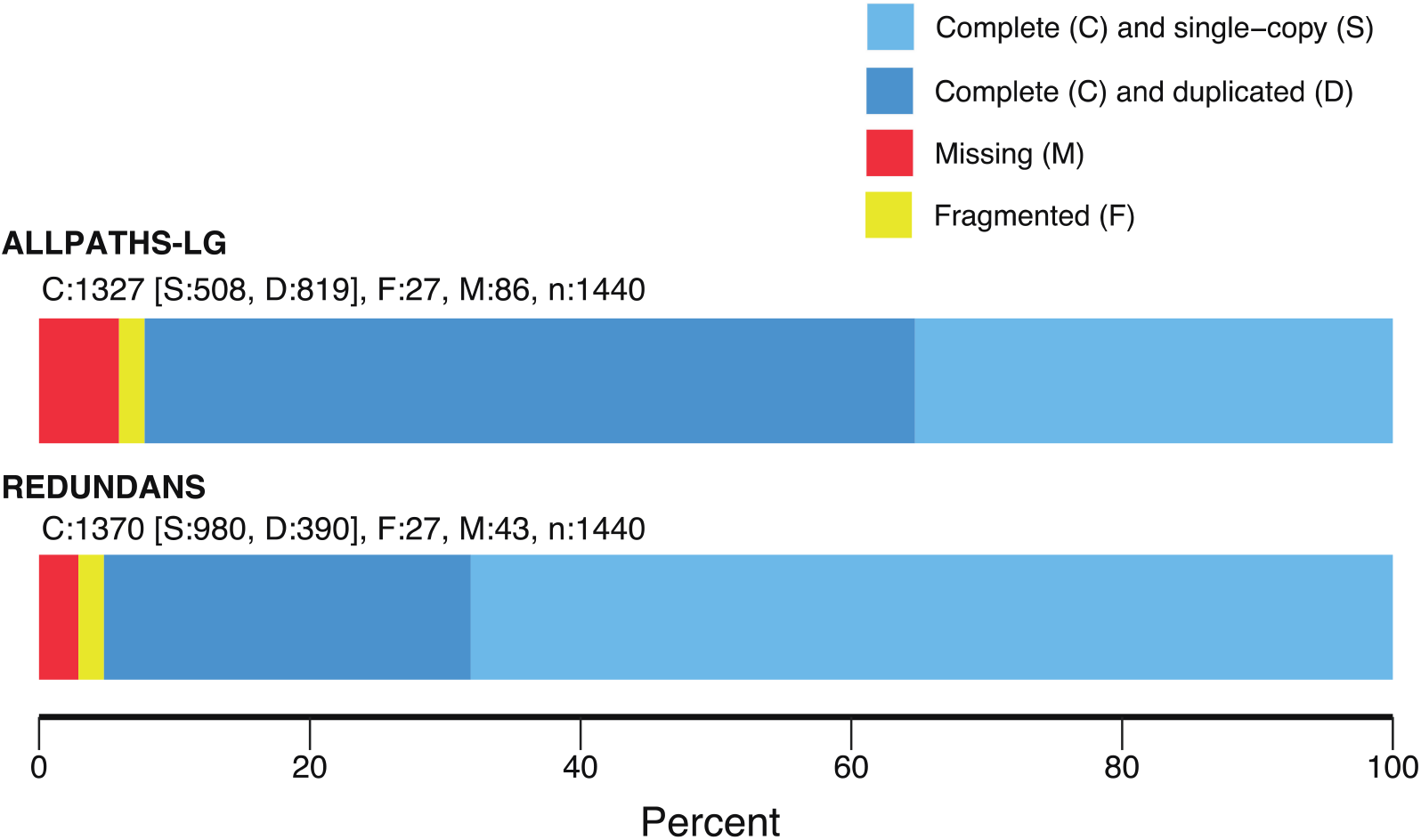
Percentage of predicted single-copy plant genes from **BUSCO** that are found one (light blue) or more times (dark blue), or are missing (red) or fragmented (yellow) in the annotated genome assembly of *Alliaria petiolata*.

Whole Genome Shotgun (WGS) sequencing produced 45.8 Gb from 229 million HiSeq reads and an additional 7.9 Gb from 15.8 million MiSeq reads. This represents an estimated 68× average coverage of the genome with an estimated genome size of 1.07 Gb. This is 80 % of the published size of 1.35 Gb, which was a rough estimate based on flow cytometry (Barow and Meister, 2002). Initial assembly with **ALLPATHS-LG** resulted in 16,743 scaffolds longer than 1 Kb. These scaffolds were linked using mate pairs. We further scaffolded by merging heterozygous loci using the **redundans** program. The final assembled genome was 1.08 Gb long across 694 scaffolds larger than 1 Kb; more than 75 % of the genome is contained in the five longest scaffolds (Table 1).

**Table 1.** Assembly statistics for the *Alliaria petiolata* genome

| Statistic | Value |
| --- | --- |
| N scaffolds (>= 1000b) | 694 |
| N scaffolds (>= 50,000b) | 227 |
| Total length | 1,075,010,735 |
| Total length (>= 50,000b) | 1,071,536,925 |
| Largest scaffold | 485,611,451 |
| GC (%) | 37.2 |
| N<sub>50</sub> | 121,941,980 |
| N<sub>75</sub> | 40,840,077 |
| L<sub>50</sub> | 2 |
| L<sub>75</sub> | 5 |
| Mean Sequence Length | 1,549,006.82 |

Cytological studies of *A. petiolata* suggest variation in chromosome number and ploidy (Cavers et al., 1979; Weiss-Schneeweiss and Schneeweiss, 2003), but are ambiguous to whether the hexaploids are allo- or auto-polyploids. Approximately 11 % of scaffolds (143 of 1,332) remained heterozygous after **redundans** assembly, compared to 36 % (6,045 of 16,743) after **ALLPATHS-LG** but prior to redundancy reduction. The average identity within these scaffolds was 99.61 %, meaning that an average of just 0.4 % of sites on the heterozygous scaffolds were still heterozygous after redundans assembly. Heterozygosity was similarly low (98.0 %) following **ALLPATHS-LG** assembly but prior to the **rendundans** step, indicating that homozygosity is a feature of the genome rather than an artifact of redundancy reduction. Finally, 27% of **BUSCO** genes (390 of 1440) were duplicated across the assembly (390 of 1440) but just 3.2 % of genes mapped to more than two scaffolds (46 of 1440), indicating that the **redundans** step was successful in collapsing homologous chromosomes.

This relatively low level of heterozygosity is consistent with a high-inbred autopolyploid or diploid. Geographic variation in ploidy (Esmailbegi et al., 2018; Weiss-Schneeweiss and Schneeweiss, 2003) with relatively simple genome make *A. petiolata* an appealing species to study the role of polyploidy in local adaptation and range expansion, which is an active area of research (e.g. (Pandit et al., 2011; Payseur and Rieseberg, 2016; te Beest et al., 2012)).

Our *de novo* transcriptome assembly included 699,048 putative isoform “transcripts” representing 535 Mb with N50 of 1,233 base pairs. The minimum transcript length was 201 as set by the Trinity default parameter while 1382 (~1.98 %) of transcripts were longer than 5 Kb. These transcripts clustered into 350,672 hypothetical genes with an average of 2.18 isoforms and 26,910 (~7.67 %) of hypothetical genes having more than five isoforms. A BLAST search of hypothetical genes to the SwissProt protein database matched 10,352 proteins with at least 90 % coverage of the query sequence, including 7,930 proteins with 100 % coverage (Table 2). The large difference in the number of protein and gene transcript is likely a function of the Trinity algorithm and the ribosome-depletion library preparation method, which includes all RNA transcripts, not just mRNA.

**Table 2.** Summary statistics of genes annotated for the *Alliaria petiolata* genome assembly.

| Statistic | Value |
| --- | --- |
| Number of genes | 64,770 |
| Number of exons | 408,155 |
| Number of introns | 343,385 |
| Overlapping genes | 9,669 |
| Contained genes | 1,464 |
| Total gene length | 210,804,785 |
| Total exon length | 102,316,121 |
| Total intron length | 109,175,434 |
| Total CDS length | 842,788 |
| % of genome covered by genes | 19.6 |
| % of genome covered by CDS | 7.8 |
| Mean mRNAs per gene | 1 |
| Mean exons per mRNA | 6 |
| Mean introns per mRNA | 5 |
| Mean gene length | 3,255 |
| Mean intron length | 318 |
| Mean exon length | 251 |
| Mean CDS length | 1301 |

Our TE annotation analysis identified 8,220 unique sequences across the *A. petiolata* genome. Of these, 112 were classified as LTRs with relatively recent origin (99 % similarity), 240 as relatively old (85 %), and 7,137 were classified as miniature inverted transposable elements (MITE). An additional 731 sequences were found to match the DNA transposase family. After masking TE sequences the final gene set from **maker** included 64,770 gene predictions with an average of 6 exons and an average exon length of 251 bp (Table 2).

A dot-plot comparison of gene synteny with the model plant *Arabidopsis thaliana* (TAIR 10) (Berardini et al., 2015) revealed large blocks of orthologous sequence (Figure 2). However, the arrangement of synteny blocks along chromosomes shows a complete re-arrangement of *A. thaliana* chromosomes when mapped to the *A. petiolata* scaffolds. Despite a high level of asynteny that is characteristic of the Brassicaceae family, the conservation of large synteny blocks holds promise for future research to identify candidate genes and genetic loci of interest for understanding plant invasions. Future research could also investigate whether ploidy variation and gene rearrangements occur among geographically and historically isolated populations of *A. petiolata*, and whether this genomic architecture has played an important role in the spread of the species.

**Figure 2.**
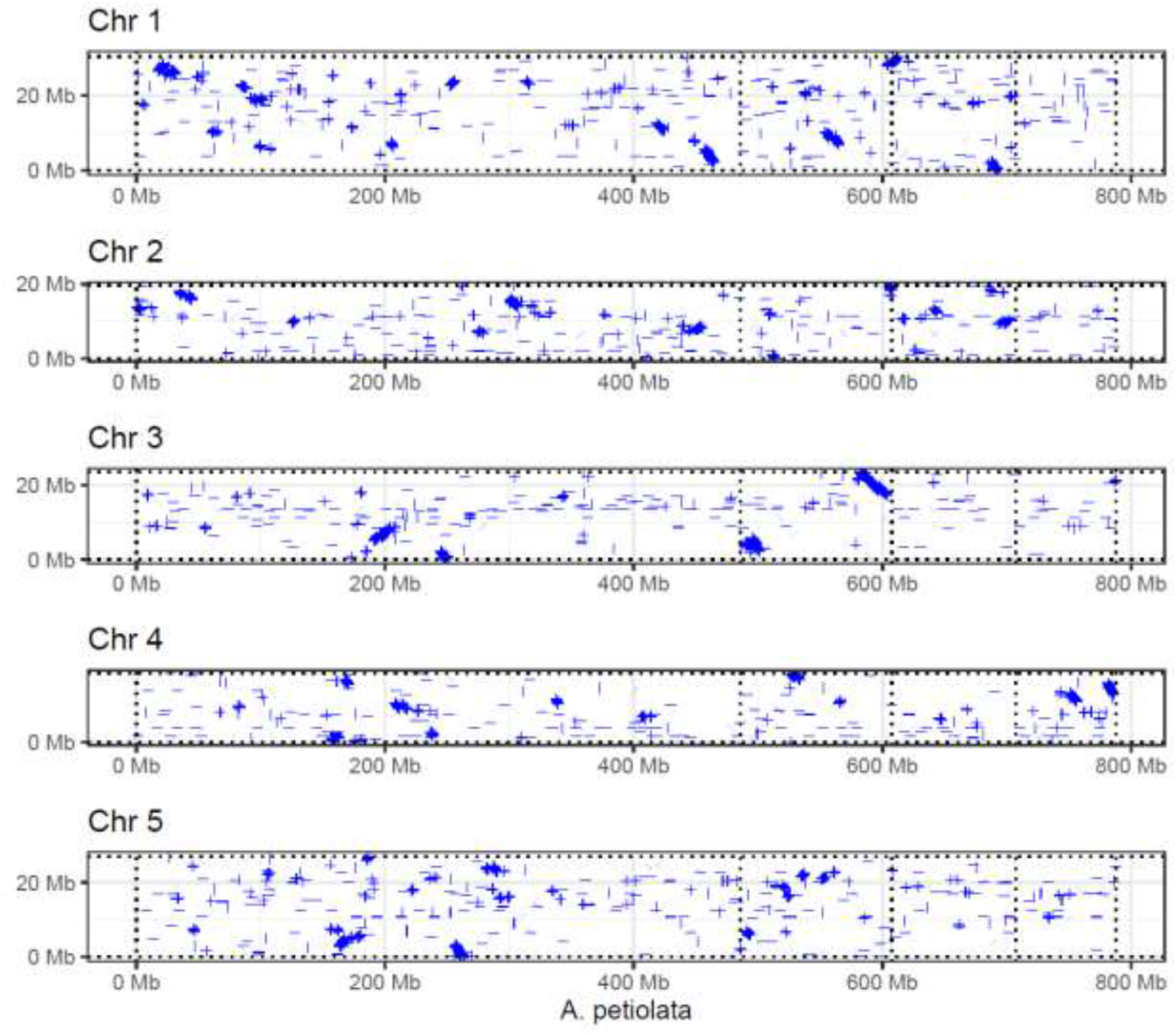
Dot-plot showing blocks of synteny between the four largest scaffolds of the *Alliaria petiolata* assembly (x-axis) and five chromosomes of the model plant *Arabidopsis thaliana*. Blue lines show aligned sequences with up to 20 % divergence. Vertical dotted lines denote separation of the major scaffolds of the *A. petiolata* assembly.

## Conclusions

There is a growing interest in the genetic causes and consequences of range expansion and biological invasion. The field of invasion genetics has emerged from ecological and evolutionary studies of invasive species but lacks well-developed model systems. The draft genome and gene annotation reported here represents an important link from the many field and experimental studies of *A. petiolata* to the genetic architecture of adaptation and invasion. High levels of self-fertility and the resultant low levels heterozygosity observed in the genome will be beneficial for future projects linking ecologically important phenotypes to specific genes. The genomic resources reported here complement available seed resources, experimental findings, and field data to accelerate genomic and molecular studies of *A. petiolata* as a candidate for a model system in invasion genetics.

## Acknowledgements

The authors are grateful for bioinformatics support from C Grassa, J Stafford, and H Schmider, hardware support from C MacPhee, and wetlab assistance from M Todesco, W Chen and A Siew. We also thank M Todesco and D Galanti for comments that improved the manuscript. This work was supported by NSERC Discovery grants to RIC and LHR.

